# The Healthy Pregnancy Research Program: Transforming Pregnancy Research Through a ResearchKit App

**DOI:** 10.1101/289371

**Authors:** Jennifer M. Radin, Steven R. Steinhubl, Andrew I. Su, Hansa Bhargava, Benjamin Greenberg, Brian M. Bot, Megan Doerr, Eric J. Topol

## Abstract

Although maternal morbidity and mortality in the U.S. is among the worst of developed countries, pregnant women have been under-represented in research studies, resulting in deficiencies in evidence-based guidance for treatment. There are over two billion smartphone users worldwide, enabling researchers to easily and cheaply conduct extremely large-scale research studies through smartphone apps, especially among pregnant women in whom app use is exceptionally high, predominantly as an information conduit. We developed the first pregnancy research app that is embedded within an existing, popular pregnancy app for self-management and education of expectant mothers. Through the large-scale and simplified collection of survey and sensor generated data via the app, we aim to improve our understanding of factors that promote a healthy pregnancy for both the mother and developing fetus. From the launch of this cohort study on March 16, 2017 through December 17, 2017, we have enrolled 2,058 pregnant women from all 50 states. Our study population is diverse geographically and demographically, and fairly representative of U.S. population averages. We have collected 14,045 individual surveys and 11,669 days of sleep, activity, blood pressure and heart rate measurements during this time. On average, women stayed engaged in the study for 59 days and 45 percent who reached their due date filled out the final outcome survey. During the first nine months, we demonstrated the potential for a smartphone-based research platform to capture an ever-expanding array of longitudinal, objective and subjective participant-generated data from a continuously growing and diverse population of pregnant women.

**Funding:** Supported in part by the National Institutes of Health (NIH)/National Center for Advancing Translational Sciences grant UL1TR001114 and a grant from the Qualcomm Foundation.

---

Historically, pregnant women, and even women of reproductive potential, have traditionally been under-represented in research, due to potential harm to the fetus.^1^ This lack of research has led to gaps in evidence-based treatment, interventions and guidelines for women,^2^ especially ones that are individualized to reflect diverse characteristics of all pregnant women.^1,3^ Since nearly four million women in the U.S. give birth every year,^4^ pregnancy is a topic that deserves far greater attention and research focus.

Given the growing population of mobile internet users which is expected to reach 60% of the global population by 2020,^5^ we have a unique opportunity to access and enroll a diverse and large population of pregnant women across the U.S. There are also a growing array of digitally connected devices and sensors, which will enable us to quantify and collect more frequently and accurately measured metrics from the expectant mother’s real world than ever before, allowing for a uniquely detailed understanding of individual variations in physiologic changes during pregnancy. In order to explore this potential, we developed a smartphone app research platform called the Healthy Pregnancy Research Program to determine its effectiveness in enrolling and engaging a large and diverse population of women throughout the duration of their pregnancy, and the ability to collect detailed and useful data that will eventually help identify the characteristics that create the healthiest pregnancy for an individual.

## RESULTS

### Recruitment and Enrollment

From the study’s launch on March 16, 2017 through December 17, 2017, there were 49,411 unique views of the welcome screen. Out of these, 9,438 self-identified that they met the inclusion criteria for the study.

As participants read through the eConsent screens, we observed a gradual drop off in the number of unique screen views. Additionally, very few people opted to read the long, detailed “learn more” version of each topic (0.4-3.8%). Interestingly, certain topics such as “Potential Risks,” and “Future Independent Research” got higher number of views for the detailed “learn more” screens (Supplementary Figure 1). The first comprehension question of the eConsent quiz was viewed by 6,558 unique users. A total of 2,186 users failed the quiz and in the end, a total of 3,777 (58% of those who started quiz) made it to the “comprehension passed” screen. However, it is unclear how many of these unique users were simply testing out the app and the eConsent process, instead of real potential users. Out of these, an additional 1,673 dropped off during the screens that gave users the option to share HealthKit data, share data with outside researchers and during the registration process that included passcode creation and email confirmation. We also filtered out users who put “Test” as their username or were clearly test users based on their name. After removing 46 individuals who were less than 18 years old in the short intake survey, we were left with a total of 2,058 participants in our study population. (Figure 1)

**Figure 1.**
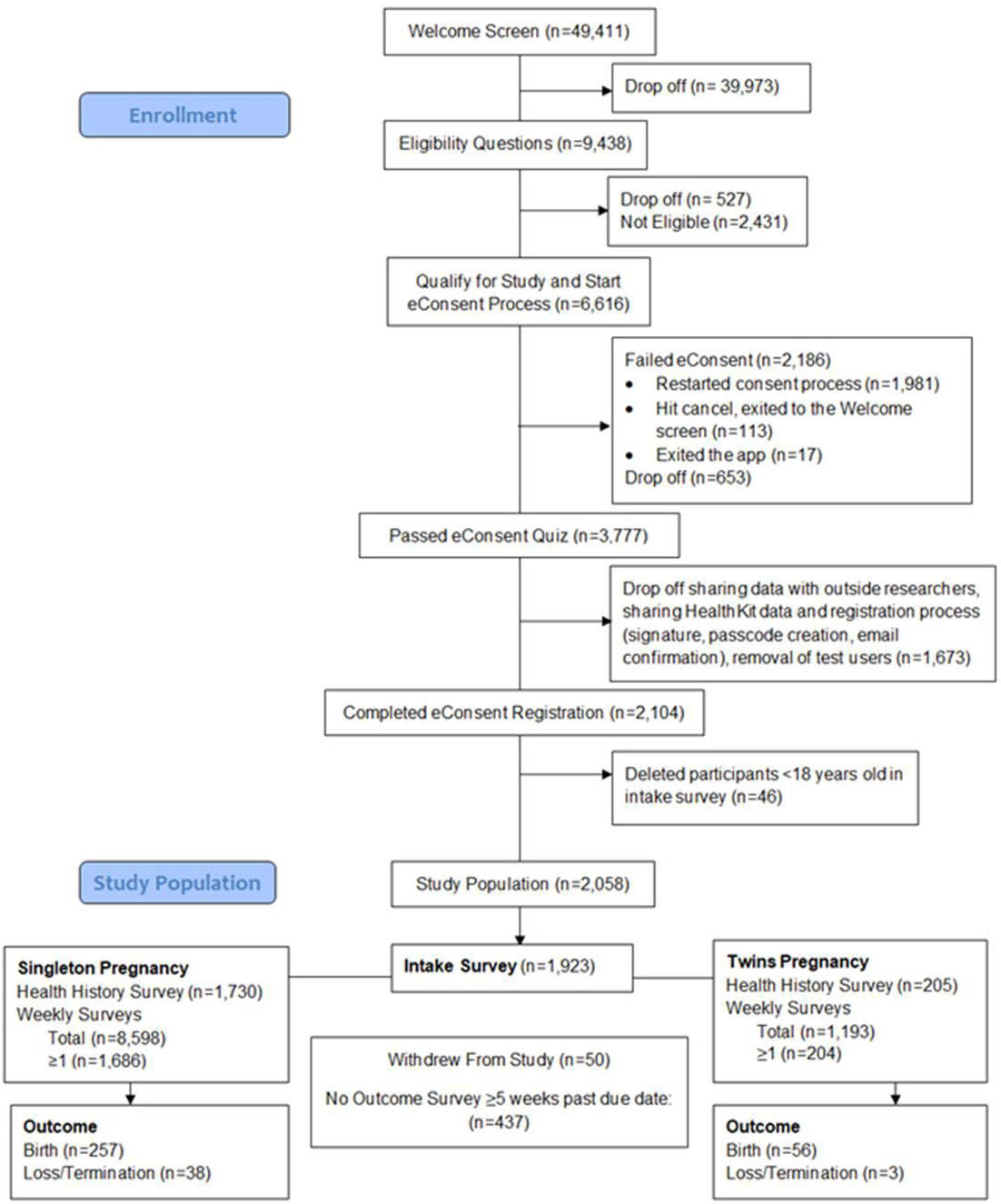
Consort diagram of participant enrollment, March 16, 2016- December 17, 2017. Participants can choose to fill out the intake survey, health history survey and weekly survey in any order.

### Sample Size and Diversity

Currently, we have enrolled 2,058 participants from all U.S. states, Puerto Rico and the U.S. Virgin Islands, with the highest number of enrollees from highly populated states such as California, Texas, and New York. (Supplementary Figure 2) Fifteen percent of our population comes from rural zip codes (zip codes without a Metropolitan Statistical Area code) compared to 19% in the entire U.S. population.^6^

We enrolled more women in the 30-39 year age group and less in the 20-29 year age group compared to national averages for pregnant women.^4^ The percentage of women in each BMI category was similar to national averages for pre-pregnancy weights.^7^ (Table 1 and Supplementary Figure 3) Although we have enrolled fewer participants from some racial minorities compared to U.S. averages, the percentage of non-White participants was very close to national averages. (Table 1)

**Table 1.** Healthy Pregnancy Population Characteristics, March 16, 2017-December 17, 2017 and US Maternal Averages.

| Characteristic | Data Missing<br>(n) | n (%) | US Average % |
| --- | --- | --- | --- |
| Short Intake Survey (n=1923) |  |  |  |
| Age (years), Mean (SD) | 35 | 30.4 ± 5.8 | 26.4<sup>*,4</sup> |
| Age Category (years) |  |  |  |
| 18-19 |  | 75 (4.0) | 4.3 |
| 20-29 |  | 788 (41.7) | 51.2 |
| 30-39 |  | 944 (50.0) | 41.5 |
| >40 |  | 81 (4.3) | 3.1<sup>4</sup> |
| Race/ Ethnicity<sup>‡</sup> | 0 |  |  |
| White |  | 1438 (74.8) | 75.7 |
| Black or African American |  | 248 (12.9) | 16.1 |
| Hispanic or Latino |  | 237 (12.3) | 23.2 |
| Asian |  | 80 (4.2) | 7.1<sup>§</sup> |
| Native Hawaiian or other Pacific Islander |  | 12 (0.6) |  |
| American Indian or Alaskan Native |  | 48 (2.5) | 1.1<sup>4</sup> |
| Middle Eastern or North African |  | 16 (0.8) |  |
| Other |  | 38 (2.0) |  |
| Live in Rural Zip Code | 76 | 280 (15.2) | 19.3<sup>6</sup> |
| Pre-Pregnancy Weight, Mean (SD) | 26 | 160.5 ± 43.4 |  |
| Pre-Pregnancy BMI (kg/m<sup>2</sup>) | 32 |  |  |
| Underweight (<18.5) |  | 105 (5.6) | 3.8 |
| Normal (18.5-24.9) |  | 850 (45.0) | 45.9 |
| Overweight (25.0-29.9) |  | 475 (25.1) | 25.6 |
| Obese (≥30.0) |  | 461 (24.4) | 24.8<sup>7</sup> |
| Share Data with All Qualified Researchers | 0 | 1404 (73.0) |  |
| Health History Survey (n=1935) |  |  |  |
| Twin Pregnancy |  | 204 (10.8)<sup>†</sup> | 3.3<sup>4</sup> |
| First Pregnancy | 13 | 629 (32.7) |  |
| Prior Miscarriage | 23 | 627 (32.8) | 16<sup>14</sup> |
| Pre-existing Conditions |  |  |  |
| Anxiety and/or Depression | 11 | 352 (18.3) | 18.1<sup>\|\|,19</sup> |
| Hypertension | 0 | 66 (3.4) | 6.8-19.0<sup>¶, 20</sup> |
| Eating Disorder | 0 | 50 (2.6) | 0.3-3.5<sup>#,21</sup> |

*SD* Standard deviation
*BMI* Body mass index
<sup>\*</sup> Age at first pregnancy, 2014
<sup>†</sup> Based on number of participants who filled out health history surveys
<sup>‡</sup> Women may identify more than one race/ethnicity
<sup>§</sup> Asian and Pacific Islander
<sup>||</sup> Percentage is for entire US population over 18 years old
<sup>¶</sup> Percentage of US women aged 20-34 years old, and 35-44 years old, respectively, in 2015
<sup>#</sup> Lifetime prevalence for US women, anorexia nervosa, (0.3), bulimia nervosa (1.5), and binge eating disorder (3.5), 2001-2003

### Engagement

We found that our study population was engaged by the study surveys with 93.4% of enrollees filling out the intake survey, 94.0% filling out the health history survey, 91.8% filling out at least one weekly survey, and 84.3% of participants opting to share HealthKit data with the study. Currently, we have collected 11,669 days of HealthKit measurements, with 11,034 days of distance run/walked measurements, 11,390 days of step measurements, 1752 days of sleep measurements, 1807 days of HR measurements, 665 days of weight measurements, and 235 days of BP measurements (Supplementary Table 1). Over the course of a single day, there were often tens to thousands of HR, distance and step count measurements from participants wearing a connected device such as an Apple Watch.

Among participants who reached their due dates, we found that participants with singleton pregnancies filled out an average of 6 surveys over 56 days (74,200 total person-days of follow up) and women with twin pregnancies filled out an average of 7 surveys over 82 days (13,202 total person-days of follow-up). Forty-five percent of women completed their final outcome survey, out of the 791 women who went to full term or had an earlier outcome (Figure 1).

### App Collected Data

During the course of pregnancy, the population average of self-reported HRs rose from 80 beats per minute (bpm) in the first trimester to 87 bpm in the third trimester. Mean HealthKit HR rose from 84 bpm in the first trimester to 92 bpm in the third trimester. Mean self-reported BP remained relatively stable from 114/71 mmHg in the first trimester to 116/72 mmHg during the last trimester, while mean HealthKit BP was higher in the first and third trimester and dropped in the second trimester. Women self-reported a weight gain of approximately 30 lbs. from their pre-pregnancy weight by the end of their pregnancy with average weights changing from 168 lbs. in the first trimester to 184 lbs. in the third trimester. HealthKit weight measurements were lower with an average of 156 lbs. in the first trimester to 179 lbs. in the third trimester. HealthKit collected sleep duration remained relatively stable during pregnancy with an average of 7 hours, while step count was highest in the second trimester and significantly lower in the third trimester compared to the prior two. A larger population size will enable us to evaluate some of these trends closer by individual characteristics. (Table 2 and Figure 2).

**Figure 2.**
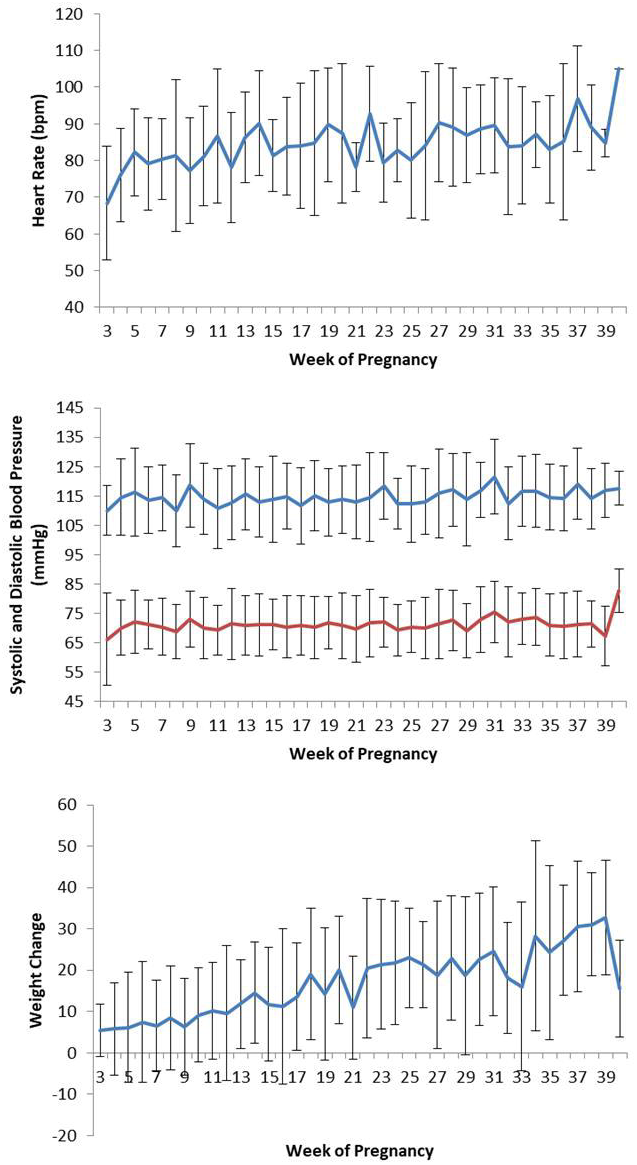
Mean (SD) physiological changes by pregnancy week from self-reported data from March 16, 2017 to December 18, 2017. (Top) heart rate (Middle) systolic (blue) and diastolic (red) blood pressure (Bottom) and weight change from pre-pregnancy weight.

**Table 2.** Population averages of physiological, sleep, and activity measurements by trimester in women with singleton pregnancies using self-reported and HealthKit collected data, March 16, 2017 to December 18, 2017

| Measurement | Trimester | Number of Measurements | Mean (95% CI) | Range |
| --- | --- | --- | --- | --- |
| Self-Reported Data |  |  |  |  |
| Heart Rate (bpm) | 1 | 1870 | 80.1 (79.4-80.7) | 24.0-153.0 |
|  | 2 | 971 | 84.8 (83.8-85.7) | 5.0-183.0 |
|  | 3 | 334 | 86.6 (84.9-88.3) | 22.0-137.0 |
| Systolic BP (mmHg) | 1 | 1799 | 114.3 (113.6-114.9) | 44.0-226.0 |
|  | 2 | 1393 | 114.4 (113.7-115.0) | 65.0-175.0 |
|  | 3 | 610 | 115.9 (115.0-116.9) | 80.0-164.0 |
| Diastolic BP (mmHg) | 1 | 1799 | 70.9 (70.4-71.3) | 7.0-123.0 |
|  | 2 | 1393 | 70.9 (70.3-71.4) | 6.0-110.0 |
|  | 3 | 610 | 72.0 (71.2-72.9) | 6.0-98.0 |
| Weight (lbs) | 1 | 3920 | 167.8 (166.4-169.1) | 70.0-356.0 |
|  | 2 | 2387 | 177.1 (175.3-178.9) | 70.0-334.0 |
|  | 3 | 860 | 183.8 (181.0-186.5) | 70.0-380.0 |
| HealthKit Data |  |  |  |  |
| Heart Rate (bpm)* | 1 | 428 | 84.0 (82.7-85.3) | 53.0-175.1 |
|  | 2 | 621 | 87.8 (86.9-88.7) | 55.5-138.5 |
|  | 3 | 506 | 91.7 (90.8-92.6) | 53.0-134.5 |
| Systolic BP (mmHg) | 1 | 49 | 120.6 (115.9-125.2) | 81.5-162.0 |
|  | 2 | 63 | 111.1 (108.2-114.0) | 88.6-139.0 |
|  | 3 | 53 | 117.3 (113.3-121.3) | 77.8-151.0 |
| Diastolic BP (mmHg) | 1 | 49 | 77.7 (74.2-81.2) | 56.0-99.0 |
|  | 2 | 63 | 70.1 (68.1-72.1) | 56.0-98.0 |
|  | 3 | 53 | 73.5 (70.7-76.3) | 50.0-93.7 |
| Weight (lbs)<sup>†</sup> | 1 | 126 | 156.3 (147.5-165.2) | 85.0-549.8 |
|  | 2 | 226 | 159.3 (155.0-163.5) | 116.4-308.0 |
|  | 3 | 206 | 179.4 (173.9-184.8) | 110.0-312.0 |
| Sleep (hours) | 1 | 345 | 6.9 (6.6-7.2) | 0.1-16.7 |
|  | 2 | 605 | 7.0 (6.8-7.2) | 0.0-16.5 |
|  | 3 | 435 | 7.0 (6.8-7.3) | 0.0-17.0 |
| Daily Number of Steps |  |  |  |  |
|  | 1 | 2624 | 2527 (2442-2611) | 1-17914 |
|  | 2 | 3718 | 2837 (2758-2916) | 4-27377 |
|  | 3 | 3123 | 2252 (2183-2320) | 6-17307 |

*CI* confidence interval
*bpm* beats per minute
\*Mean daily heart rate
†If multiple weights were taken per day, first weight of the day was used.

The top ten most prescribed medications from the health history survey included drugs to treat or prevent anti-depression, thyroid deficiency, miscarriage, morning sickness, and diabetes. The top ten most taken over the counter drugs were prenatal vitamins, and drugs for pain relief, allergy, nausea and heartburn, sleep, and probiotics. Although, the FDA is moving away from letter risk categorization of medications, 6 of the top 10 prescribed medications and 2 of the top 10 over-the-counter drugs were category C, meaning risk is not ruled out and animal reproduction studies have shown possible adverse effects in the fetus (Table 3).

**Table 3.** List of the top ten most prescribed and over-the-counter medications during pregnancy, drug indication, FDA pregnancy category, and percentage of participants taking the drug. Data is from the health history survey for singleton pregnancies, March 16-December 17, 2017, n=1,730

| Prescribed Medications | Over-the-counter |
| --- | --- |
| 1. Antidepressants (8%) | 1. Prenatal Vitamins (93%) |
| a. Zoloft, C (4%) | 2. Analgesic (9%) |
| b. Bupropion, C (1%) | a. Acetaminophen, B (5%) |
| c. Prozac, C (1%) | b. Aspirin (NSAID), C/D (4%) |
| d. Celexa, C (1%) | 3. Allergy (7%) |
| e. Lexapro, C (1%) | a. Zyrtec, B (3%) |
| 2. Levothyroxine (Thyroid Deficiency), C (5%) | b. Claritin, B (2%) |
| 3. Progesterone (Infertility/Prevent Miscarriage), A (2%) | c. Benadryl, B (2%) |
| 4. Morning Sickness (3%) | 4. Zantac (Nausea/Heartburn), B (2%) |
| a. Diclegis, A (2%) | 5. Unisom (Sleep), B (2%) |
| b. Zofran, B (1%) | 6. Probiotic (Healthy Gut), Not Assigned (2%) |
| 5. Metformin (Type 2 Diabetes), B (1%) | 7. Tums (Antacid), C (1%) |

## DISCUSSION

During the first nine months of the deployment of the Healthy Pregnancy ResearchKit app, we have enrolled over 2,000 participants from 50 states, and collected over 14,000 individual surveys and 11,000 days of HealthKit measurements. Going forward, as we incorporate a greater number of enrollees, we plan to incorporate more novel digital devices for an even greater variety and volume of participant-generated data. Overall, we believe this app can prove to be on ongoing, ever-improving source of important insights to better understand the individual factors that create a healthy pregnancy for all women.

Over the last decade there has been a shift in how pregnant women find and share pregnancy related health information. A recent study found through a survey of pregnant women that 55% were using an app related to pregnancy, birth, and/or child care.^8^ In fact, 7% of all mHealth apps in 2015 were focused on women’s health and pregnancy.^9^ These apps provide useful tailored information based on week of pregnancy at their fingertips. High pregnancy app usage combined with an increasing number of people using digitally connected devices and sensors to monitor their health, allows for unique possibilities to collect and provide information that is tailored to individual consumers. These technologies also have the ability to transform research by enabling the sharing of participant-generated data in a way that is less burdensome to participants.

One unique aspect of the Healthy Pregnancy Research Program is that it is embedded in WebMD’s highly trafficked pregnancy app (Figure 3). Since 2013, over 1.6 million people have downloaded WebMD’s Pregnancy App. WebMD’s Pregnancy App is a leader in providing medical related information to pregnant women, is medically reviewed by physicians and provides answers to many questions that women would typically ask their health care provider. We believe that by incorporating our study platform into a trusted app where pregnant women are already going for their information has given us high visibility and likely better long-term engagement. Efforts are underway to embed our app into other trusted digital platforms.

**Figure 3.**
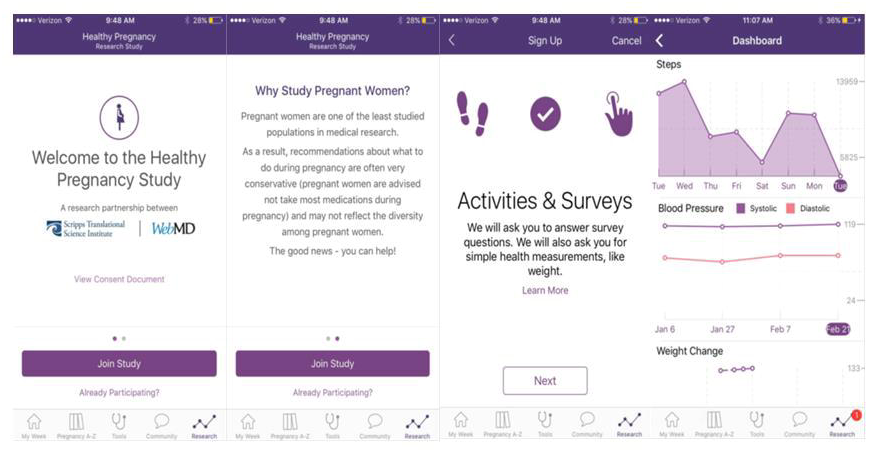
Screenshots of the Healthy Pregnancy Study. Left to right: (1) Welcome screen (2) Why study pregnant women? (3) Activities and surveys (4) Dashboard tracking physiological and activity changes during pregnancy.

We found that participants were more likely to read the detailed version of specific eConsent topics (e.g. Your Privacy, Potential Risks, and Future Independent Research), which may indicate additional interest, confusion, or concern for these topics. Currently we observed substantial drop-offs throughout the eConsent process, with 33% of participants failing the short eConsent quiz. In the future, giving participants who failed an eConsent question a description of why their answer was incorrect, instead of forcing them to re-do the entire eConsent process, in other words, restructuring the quiz from a summative to a formative evaluation, may improve enrollment. This teaching/reinforcing approach may also help improve participant understanding within the consent process. Additionally, identifying which questions were answered incorrectly is important for improving our understanding of where we are failing to adequately explain topics to participants and perhaps explore additional formats for eConsent presentation, such as voiceover or video.

Relative to other ResearchKit studies, we have seen less attrition and a higher percentage or participants filling out the surveys.^10–13^ Additionally, 73% of participants in our study were willing to share data with outside researchers which is similar to other ResearchKit studies (67-78%).^10,12,13^ Higher engagement and willingness to share data may be a result of our unique study population of expectant mothers bringing their baby into the world, during a limited, circumscribed temporal window. We may also have lower drop-off because women receive other useful information through the WebMD pregnancy app and may therefore be less likely to delete it and withdraw from the study. In order to improve long-term engagement, it will be necessary to continuously refine the return of useful information to participants. As more data is gathered from a greater number of women it will become possible to eventually match a woman with their “digital twin” to help more precisely guide their expectations and health behaviors.

Certain participant characteristics seem to influence their likelihood of participating. Interestingly, 3% of all births in the U.S. are twin births;^4^ however our study is capturing a much larger proportion of twin pregnancies (11%). Similarly, we saw a higher percentage of women joining our study with prior miscarriages (33%) compared to national averages (10-20%).^14^ (Table 1) Women also joined our study on average at gestational week 17. Finding ways to encourage women to join earlier will allow us to collect important data that may help us identify signs of early complications. Overall, higher engagement among certain sub-populations or during certain times of pregnancy may be a result of heightened concern or increased questions during these times or for these groups. These groups may be more motivated to be part of a research study with the hope of answering their questions or improving their particular condition.

Traditional clinic based studies have often lacked participant diversity due to participation challenges such as lack of access to care centers that typically refer people to research studies, time constraints for participation, difficulties with transportation to research sites, lack of childcare while participating, and limited participation hours.^15^ By conducting an app-based research study that takes limited time, and can be done anywhere and at any time, we have helped overcome some of these challenges, which has likely increased our participant diversity. Future expansion to Android phones is also essential and will enable us to access a much wider and varied user population in the U.S. and eventually, globally.

As the availability of an increasing variety of wireless, connected sensors grows, we anticipate including the automated daily (or even more frequent) collection of multiple parameters known to be germane to pregnancy such as BP, HR, activity, sleep, stress, nutrition, and glucose levels. We can also assess the impact of new digital platforms and home-based sensors at improving positive behavior change to improve health. In addition to conducting research, a primary future objective of this study is to help women meaningfully interpret and understand their personal data through visualizations, risk profiles, and comparisons to other individuals like them. Ultimately, this will make for more informed decisions for pregnant women when it comes to things from medication choices, to healthy weight gain and ideal sleep during pregnancy.

In summary, we aim to use our app-based research platform to help fill in many knowledge gaps that exist for pregnancy, ranging from individualized weight gain recommendations to earlier identification of preeclampsia, gestational diabetes, peripartum depression, and other complications. The first nine months of our study have proven the app’s success to quickly and easily collect detailed and useful data remotely from a large and relatively diverse population from across the U.S. These data may inform both improved population based public health interventions and individualized care that will ultimately help create healthier pregnancies.

## METHODS

### Overview and Goals of the Research Program

The Healthy Pregnancy Research Program is a ResearchKit app-mediated study developed by Scripps Translational Science Institute (STSI) and WebMD. ResearchKit is an open-source framework for building research apps, created by Apple.^16^ It allows researchers to conduct studies quickly, cheaply, and easily by collecting survey and HealthKit data through a dedicated study app on the participant’s iPhone.^16,17^ Our ResearchKit study can be accessed via the WebMD Pregnancy app (Figure 3) or can be downloaded directly from the iTunes store.^18^

Data collected include maternal weight change, blood pressure (BP), medication usage, symptoms, diagnoses such as preeclampsia and gestational diabetes, as well as birth outcomes, primarily through participant-reported surveys. We are additionally collecting maternal activity data (steps and distance run/walked), sleep duration, weight, HR (heart rate), and BP directly through the Apple iPhone’s HealthKit. The majority of HealthKit data is likely automatically uploaded from sensors; however, it is possible that some of the HealthKit data is also self-reported and manually entered. Currently, activity, weight and BP data is plotted in a graph so that participants can see how their individual data changes throughout their pregnancy.

This study was approved by the Scripps Research’s Institute’s Institutional Review Board on February 17, 2017. The study was also registered on ClinicalTrials.gov (Identifier: NCT03085875) on March 21, 2017.

### Qualification and eConsent

Any pregnant person, 18 years or older, who lives in the U.S. and is comfortable reading and writing on their iPhone in English is eligible to join our study. After self-identifying that they qualify for the study, potential participants self-navigate an electronic Consent process (eConsent) that includes 17 screens highlighting key consent topics ranging from data sharing and data privacy, to potential risks. The consent process is entirely self-guided and self-administered, with no in-person steps. Within the eConsent process, participants can click on a “learn more” option to get a more detailed description of each topic or advance to the next screen. If a potential participant has questions regarding the eConsent, they are given the contact information of the study coordinator and IRB. After completing the eConsent, participants must correctly answer four questions that test their comprehension of core study topics in order to join the study. Those who do not answer correctly are directed to the beginning of the eConsent process.

### Surveys

After completing the eConsent process, participants are given a short intake-survey, health history survey and a weekly survey. The weekly survey recurs every week and initially asks participants if they are still pregnant, had a miscarriage or stillbirth, if the pregnancy ended, or if they gave birth. If they are still pregnant, the weekly survey asks recurring questions about physiological measurements, medications, and symptoms. If a participant had a prenatal visit during that week, they are given a few additional questions about vaccines and diagnoses. If they indicate a birth they are given the outcome survey questions about the baby’s weight and length, and labor and delivery questions. An outcome survey is also given 4 weeks after a participant’s due date, if they don’t indicate a birth outcome during the weekly survey. At any time, participants are able to skip over questions that they do not want to answer. (Supplementary Table 2)

## DATA AVAILABILITY

The datasets generation during and/or analyzed in this study are available from on reasonable request.

## ACKNOWLEDGMENTS

We would like to thank the work of our study manager, Gail Ebner and our study coordinator, Lauren Ariniello. We would also like to thank Michael Galarnyk for his help summarizing the HealthKit data. Finally, we would like to thank all the developers at WebMD for their work with the app.

## CONTRIBUTIONS

Guarantor: Radin had full access to the study data and takes responsibility for the integrity of the complete work and the final decision to submit the manuscript. Study concept and design: Radin, Steinhubl, Su. Acquisition, analysis, or interpretation of data: All. Drafting of the manuscript: Radin, Steinhubl. Critical revision of the manuscript: All. Obtaining funding: Steinhubl, Topol, Greenberg.

